# Temporal changes in macrofungal alpha diversity over four decades in Europe

**DOI:** 10.1101/2025.11.16.688674

**Authors:** Franz-Sebastian Krah, Claus Bässler, Max Zibold, Jacob Heilmann-Clausen

## Abstract

Terrestrial organisms have shown mainly positive or neutral changes in local richness across late-and post-industrial decades. A lack of time-series data has so far hindered the exploration of similar changes in fungal alpha diversity. Here, we analysed 5 million macrofungal records and tracked changes in alpha diversity between the 1980s and 2010s across 500 time comparisons. Using a novel tool to standardize for geographic spatiotemporal sampling effort, we estimated alpha diversity using coverage-based rarefaction and extrapolation. We found weak positive net changes in alpha diversity across most of Europe. Drier conditions and nitrogen increases were linked to fungal alpha diversity decreases, while climate warming increased alpha diversity across fungal functional groups. Our results suggest that local-scale factors, such as drought and chemical pollution, are more constraining to fungal biodiversity than regional-scale climate warming and habitat loss.

## Introduction

European ecosystems are significantly affected by global change, including climate and land-use change, habitat fragmentation, pollution, and invasive species ^1^. A key question in ecology and conservation is whether these changes affect biodiversity. Large-scale studies have found weak or non-significant local richness changes in plants ^2–4^, and a recent meta-analysis found mainly neutral or positive richness changes across several terrestrial organism groups ^5^, however, the kingdom of fungi was missing. Fungi constitute a mega-diverse kingdom, and species diversity is tremendously rich, with more than 10,000 macrofungal species in Europe alone ^6^. Fungi perform important ecosystem processes ^7^, such as decomposition ^8^ and the formation of mycorrhizal symbiosis, thereby substantially supporting primary production^9,10^. Thus, changes in fungal diversity may significantly affect ecosystem functioning. While several spatial studies have demonstrated that fungal diversity is strongly linked to climate across large spatial scales, followed by soil and vegetation ^11–15^, the response of fungal diversity to climate change is mainly known from a limited set of local survey studies ^16,17^. Despite the importance of space-for-time approaches, inferred interpretations on temporal changes related to climate change are uncertain ^18,19^ and may lead to erroneous conclusions ^20^. Further, other global change drivers, e.g., pollution or land-use change, have rarely been included in large-scale studies of fungal diversity, and may have overriding effects ^21^.

The few available fungal time series are extremely local (plot-level) and have found partly increasing or decreasing richness across 1975 to 2006 and 1995 to 2013, respectively ^16,17^. Due to the sharp scarcity of replicated time series in fungi, we lack knowledge on how their diversity changed across past decades on the landscape to the continental scale. Despite the crucial roles fungi play in terrestrial ecosystems, their alpha diversity response to global change in the past decades remains unknown. Our overall aim with this study is to shorten this knowledge gap, focusing on macrofungal alpha diversity changes in Europe over the last four decades. In line with studies focusing on other terrestrial organisms ^2,3,5^, we expect limited net diversity change. Further, analyses of spatial patterns of fungal alpha diversity at the continental to global scale have found that fungal richness generally showed a hump-shaped relation with mean annual temperature ^15,22^, suggesting that fungal richness can be assumed to increase with climate warming in colder parts of Europe, while potentially decreasing in warm zones. Spatial studies have also found positive responses to precipitation, especially in saprotrophic basidiomycetes ^15^. A large-scale drought experiment revealed a predominantly high degree of drought resistance and resilience in fungal diversity across Europe ^23^. We thus expect no significant changes to precipitation or moisture. Further, a recent review of soil fungi has highlighted the role of nitrogen deposition with mixed directional effects depending on fungal functional groups ^24^. We expect negative effects of nitrogen increase on ectomycorrhizal fungi as this lifestyle has been negatively affected by nitrogen addition ^25^. Aside from these concrete hypotheses, we also explore other environmental factors relevant to the local-to landscape-scale. We categorized our predictor variables into base and change predictors reflecting macroenvironmental (macroclimate, human footprint index), microenvironmental (microclimatic temperature and light conditions), and edaphic (moisture, nitrogen, pH) predictors. The change predictors reflect anthropogenic changes, e.g., temperature increase through climate change ^26^, changes in light conditions through forest management ^27^, and nitrogen increases through nitrogen deposition and fertilization ^28^.

To test our hypotheses, we compiled a multisourced European-scale fungal fruiting dataset including all Agaricomycetes, covering the 1970-2024 and 25 million records, which we assembled into 50km x 50km grid cells. After data cleaning, 5 million records with 9.000 species were analysed. Diversity change was quantified between 1970-1990 and 2014-2024, two time intervals selected to increase diversity estimation accuracy and temporal coverage. Microenvironmental and edaphic factors were calculated based on plant Ellenberg indicator values for 2,500 herbaceous plant species within the same grid cells as the fungal data. Apart from total macrofungal diversity, we considered two functional groups of fungi, namely ectomycorrhizal fungi, which form symbiosis with woody plants, and free-living saprotrophs, which degrade dead organic matter ^29^. We estimated richness using the recently developed Hill number–based methods ^30,31^ for quantifying alpha diversity, standardized by sample coverage, providing a powerful framework to correct biases from spatiotemporally imbalanced sampling and distinguish responses of rare and dominant species along the Hill series.

## Material and Methods

### Fruiting and climate data

We downloaded all fungal sporocarp records from the Global Biodiversity Information Facility (GBIF) with the query term ‘Agaricomycetes’. This resulted in 19,225,366 records (17 January 2025, dataset, https://doi.org/10.15468/dl.j6m9cz). We added 1,043,262 records from the French National Fungal Database ^32^ and 4,560,050 records from the German Mycological Society. We thus started with 24,828,678 records. We processed the data in the following steps: (1) exclusion of all records without coordinate data, (2) exclusion of all records without exact year information, (3) exclusion of all records before the year 1970 and after the year 2024, (4) exclusion of all records with a coordinate accuracy worse than 10 km, (5) exclusion of taxa resolved above the species level (e.g. genera, family, order), (6) exclusion of records not based on fruiting bodies (e.g. eDNA from metabarcoding projects), (7) harmonization of the data from different sources by updating the taxonomy (using the GBIF taxonomic backbone), by removing duplicate records (base on species, coordinates and date), and by excluding all taxa from the German and French inventories other than ‘Agaricomycetes’, (8) exclusion of flagged records using the R package CoordinateCleaner, developed for cleaning GBIF data ^33^, (9) assignment of each record to a calendar week (12) exclusion of all records recorded outside Europe (10) assignment of each record to a 50km x 50km grid cell.

For the analysis, we aggregated the data into spatial and temporal units and added environmental variables. Specifically, we used a 50km x 50km grid cell because it is the most widely used grain size for fungal macroecology ^11,13,34^ and balances data sufficiency and environmental heterogeneity. We further separated the data into a base survey and resurvey time interval, namely 1970-1990 and 2014-2014. We chose to use these time windows to gain meaningful and robust assemblage data in both time intervals. Our data stems from unstandardized collections, with a sharply increasing trend. Hence, we used a 20 year time span for the baseline time window compared to the recent (1970-1990: 1,058,004 vs. 2014-2024: 4,255,173 records).

Finally, we separated the dataset into a dataset with all species and two datasets for ectomycorrhizal and saprotroph species, respectively, based on Ref^35^. The trait database contains primary lifestyle coding, where we used “ectomycorrhizal” directly. Saprotrophs on the other hand were aggregated from the lifestyle categories “soil_saprotrophs”, “litter_saprotrophs”, “wood_saprotrophs”, “dung_saprotrophs” and “unspecified_saprotroph”.

### Diversity estimation

Occurrence records are often incomplete and affected by various biases. Because most of the analyzed records originate from activities not specifically designed to assess diversity patterns, their spatial coverage is typically uneven and unsystematic, leading to substantial omission errors ^36^. To reliably detect patterns and changes in species diversity, it is essential to account for geographic variation in sampling completeness. Using sample coverage enables the standardization of observed diversity through rarefaction or extrapolation based on sample completeness ^30^. This approach allows meaningful comparisons of species diversity within and among assemblages with differing sampling intensity in space and time, even without knowing the true underlying diversity ^36^. Coverage-based rarefaction and extrapolation mitigate biases related to uneven sampling activity. Sample coverage is estimated based on the proportion of singletons. Diversity is rarefied and extrapolated to a given level of sample coverage. Coverage-based standardization is a mathematically elegant and statistically robust way to standardize samples and has been increasingly used in the ecological literature (e.g., Ref. ^37^), and for occurrence records data (e.g., Ref. ^38^). We therefore used standardization by the sample coverage across time intervals within each grid cell.

Our data contained the number of unique records of a species within a 50km x 50km grid cell. However, the number of records is an unreliable measure of abundance ^39^. Therefore, we used an approach based on the incidences of species within the grid, i.e., the number of subgrids occupied, as proposed by Ref^38^. We first prepared a spatial grid with 50km x 50km grid cells using the ETRS89 coordinate reference system (ETRS89 = European Terrestrial Reference System 1989 + Lambert azimuthal equal area (LAEA) projection (EPSG: 3035)), which has the advantage that grid cells are equally sized across the spatial gradient and thus provide the same area-based estimates for diversity and environmental variables. Each 50km x 50km grid cell was divided into 25 subgrids measuring 10km x 10km for each time interval. We quantified the frequencies of species within the subgrids as presence and absence. This approach thus contained an incidence matrix for each grid (a species x subgrid matrix with 0s and 1s), which we used for coverage-based alpha diversity estimation over time for each grid cell. For alll sample-coverage based calculations we used the *iNEXT* R package ^40^, specifically the functions *DataInfo* and *estimateD* and used both time interval incidence raw matrices as input (list).

Given the known limitations of diversity estimation methods, we excluded grid cells with insufficient sampling. Specifically, a grid cell was removed if it contained fewer than 50 observed species (25 for lifestyles). The remaining grids had an average sample coverage of 0.63 and 0.85 for the first and second time intervals, respectively. We then used the function estimateD for every grid using the lower observed sample coverage at double sample size. Based on the estimated diversity, we computed the diversity change between the second (resurvey) and first (baseline survey) in each grid as the natural log response ratio, proposed in Ref^4^. We calculated the changes in alpha diversity as the natural log response ratio per time interval as:

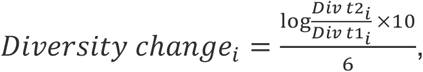

where “Diversity change_i_” is the change in diversity of grid i; Divt1_i_ and Divt2_i_ are respectively the diversity at the resurvey and baseline year in grid i. We used 6 as a denominator, indicating the 6 decades the datasets are maximally apart. Note this choice does not change the results. We chose 6 to reach magnitudes of change comparable with previous studies. Log ratios express proportional change, making changes between grid cells with different total diversity levels comparable. Since all time series were equally long, it is a constant. To test if the level of sample coverage used for rarefaction and extrapolation had an effect on the diversity change, we explored their relation. We found a net negative diversity change below a sample coverage of 0.5 and a net diversity change of 0 above 0.5 (Fig. S1A). We therefore excluded grids with sample coverage below 0.5 as they were biased towards negative diversity change. To further test if chosing a different sample coverage level per grid affected our results we also used a predefined level of 0.5 across grids. We found a strong correlation (r>0.95) between both (Fig. S1B). We therefore used diversity change based on sample coverage uniquely set for each grid to maximise data usage per grid. Using this approach, we obtained 521, 433, and 480 grid cells temporal comparisons for all fungi, ectomycorrhizal fungi, and saprotrophic fungi, respectively.

We quantified taxonomic diversity using Hill numbers at orders *q* = 0, 1, and 2, which account for differences in relative abundances. The interpretation of Hill numbers is as follows: for *q* = 0, all species contribute equally, regardless of their frequency, making this measure particularly sensitive to rare species. For *q* = 1, species are weighted according to their relative frequencies, thus reflecting the diversity of common species. For *q* = 2, frequent species are given disproportionately higher weight, so the resulting measure primarily reflects the effective number of dominant species. For alpha diversity, q = 0 corresponds to richness, q = 1 to Simpson index and q = 2 to Shannon index.

### Environmental data

We chose a set of environmental variables that reflect macroenvironmental broad regional-scale effects (macroclimate, human footprint index). The local-scale conditions we separated into microenvironmental effects (temperature and light) and edaphic effects (nitrogen, moisture, and reaction (pH)).

Macroclimatic variables were computed based on the CRU 4.09 dataset ^41^. We calculated temperature and precipitation based on averages and sums for the months March to October, which better reflect the physiological activity of fungal mycelium and fruiting. Please note that we also explored March-December and May-December, and the base and change variables showed correlation coefficients above 0.99. For climatic variables, we calculated the average for the base time interval (t1: 1970-1990) and for the end time interval (t2: 2010-2024), and the change in mean as t2–t1. For the human footprint index (HFI), we utilized gridded terrestrial data available yearly between the earliest year, 2000, and 2018 ^42^ with 2000 as the base year and 2018 as the end year. The human footprint index includes eight variables (built environment, population density, nighttime lights, cropland, pasture, roads, railways, and navigable waterways) ^42^ and thus reflects mainly habitat loss in comparison to the other variables selected which focus on climate and edaphic changes through human impacts.

For the microenvironmental and edaphic conditions of fungal habitats, we calculated the plant-community average Ellenberg indicator values for each grid cell and their changes based on herbaceous plant species ^43^. We first downloaded records from GBIF with the search term “Magnoliopsida” and restricted the search to a coordinate uncertainty below 1000m, records with coordinates, and Europe as a continent. We repeated this process for the same time intervals as those used for the fungal data (1970-1990 and 2014-2024). We then used coordinateCleaner with additional cleaning steps as above to remove spurious coordinates. We then prepared a community matrix with grid cells and the subset of species with Ellenberg indicator values. We restricted the plant species to herbaceous plants using the woodiness database ^44^. This resulted in 2485 herbaceous plant species for both time intervals with sufficient data. Since records do not reflect abundances, we used a presence/absence (1/0) coding and computed the community mean indicators Moisture (M), Reaction (R), Nitrogen (N), Light (L) and Temperature (T). For maps of the predictor values, see Fig. S2. They align well with expectations of their intensity across Europe. We used the first time interval as base and the difference of t2-t1 as change. Since the base and change values were correlated (r>0.7, Fig. S3) we calculated the residuals of change against base, which reflects the removal or addition of the value compared to the base. We gridded all environmental variables on the ETRS89 coordinate reference system. We applied tangent, logarithm, square root, or arc-hyperbolic-sine (asinh) transformation to reach normal distribution or suppress extreme central values. We further added the biogeographic region for the grid cells based on Ref^45^.

### Statistical analysis

We were first interested in the net change of alpha diversity across Europe. We used histograms to show the distribution of diversity change for the functional groups and Hill numbers. We show the zero value and mean as vertical lines in the histograms. We further tested whether the distribution differed from zero using one parametric T-test (function t.test), and two non-parametric tests, namely the Wilcox rank sum test (function wilcox.test) and the Dependent-samples Sign-Test (SIGN.test from BSDA package). We only considered the mean of the distribution as different from zero if all three tests showed a significant deviation.

We were secondly interested in the effects of base and change environmental effects on diversity change. Using both the base and the change residuals as predictors in the same model violates basic statistical assumptions because the base and the change variables are not independent statistically (residuals). Therefore, we use two models. The “base model” with base predictors and the “change model” with the change predictors. Within both models, we selected predictors that showed a maximum correlation coefficient of 0.7 ^46^. The selected predictors for the base and change models are shown as a correlation matrix in Fig. S3. We first used linear models and tested for spatial autocorrelation within the residuals using Moran’s I and found significant spatial autocorrelation and values of ca. 0.3. We thus used generalized additive models (GAMs) that included base or change values as parametric predictors and added a spatial term, “+ s(X, Y, bs = “gp”)”. We chose the low rank Gaussian process smooth term because it accounts for distance-based similarity between nearby sites while avoiding biased inference from unmodelled spatial structure ^47^. By including this spatial term, the signal in the residuals was reduced to a Moran’s I below 0.1. Since we used the response variable within the base and change model, we used Bonferroni adjustment of the p values by multiplying them by 2. We fitted the same model separately for the Hill numbers (q = 0, 1, 2) and functional groups and display results as forest plots with estimates and their 95% confidence intervals. We also present scatter plots for all predictors, functional groups and Hill numbers along with the partial regression line form the GAM models in the Supplementary Material. Based on the fitted GAM models, we also tested the relative effect of three predictor sets using hierarchical partitioning using the function gam.hp from the gam.hp R package ^48^. As predictor sets within the base model, we used macroenvironment (mean annual temperature, precipitation sum), microenvironment (Ellenberg T and L), and edaphic (Ellenberg M, N, R). As predictor sets within the change model, we used macroenvironment (mean annual temperature change, precipitation sum change, Human footprint index change), microenvironment (Ellenberg T and L change residuals), and edaphic (Ellenberg M, N, R change residuals). We finally explored diversity change within different biogeographical regions and displayed the scaled diversity change on maps.

## Results

Fungal alpha diversity mainly showed non-significant changes over time across Europe (Fig. 1). For all fungi, richness (q = 0) showed a slight significant increase, while Shannon (q = 1), and Simpson diversity (q = 2) showed net changes not different from zero (Fig. S4). For ectomycorrhizal and saprotrophic fungi, richness and Shannon diversity showed no significant net trends, while Simpson diversity showed a slight but significant positive increase (Fig. 1A). We further explored the net diversity change in five biogeographical regions. We found a central tendency around zero, which was mostly consistent with the overall results (Fig. S5). However, the Atlantic region showed a substantial negative diversity change (Fig. S5). Based on the mapped diversity change, diversity loss was most pronounced in the United Kingdom, and diversity gain was associated with northern boreal areas (Fig. 1). This pattern was consistent across functional groups.

**Fig. 1.**
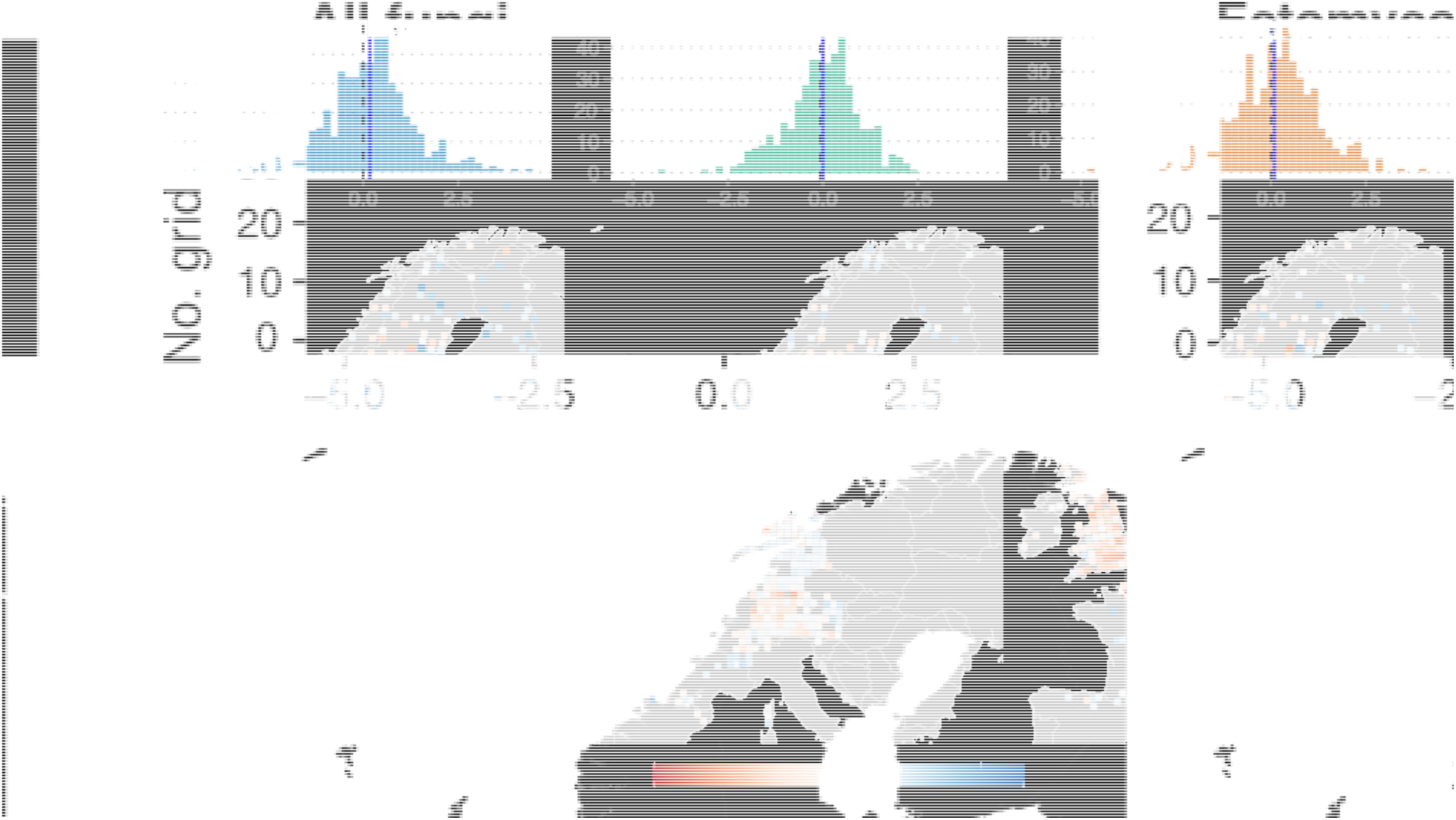
Changes in alpha diversity across decades in European fungal fruiting communities. Alpha diversity is based on a coverage-based rarefaction and interpolation of incidence data per grid cell. The diversity change was calculated as log ratios between 1970-1990 and 2014-2024. Histograms of diversity change for richness (q = 0, rare). The zero point and change mean are drawn as dashed and solid lines, respectively. The mean was tested against zero using one parametric and two non-parametric tests, and we considered their significance (indicated by a star) only if all three tests showed a p-value of less than 0.05. Maps showing the rescaled richness change (q = 0, rare).

To gain an understanding of the links between base and change predictors and fungal diversity change, we employed generalized additive models that incorporated macroenvironmental, microenvironmental, and edaphic predictors and a spatial autocorrelation term. We found that diversity change responses were overall equally well explained by the predictors in the base and change model, with a mean of 22% explained deviance. Using hierarchical partitioning, we found that diversity change responses were mainly explained by macro-and microenvironmental predictors in the base model. In the change model, diversity changes were, in contrast, mainly explained by macroenvironmental and edaphic predictors. For ectomycorrhizal fungal diversity, the relative importance of predictor sets largely followed the response of all fungi. Saprotrophic fungal diversity was more strongly explained by edaphic predictors in the base model. In the change model, saprotroph diversity change was also mainly explained by macroenvironmental and edaphic factors (Fig. 2A).

**Fig. 2.**
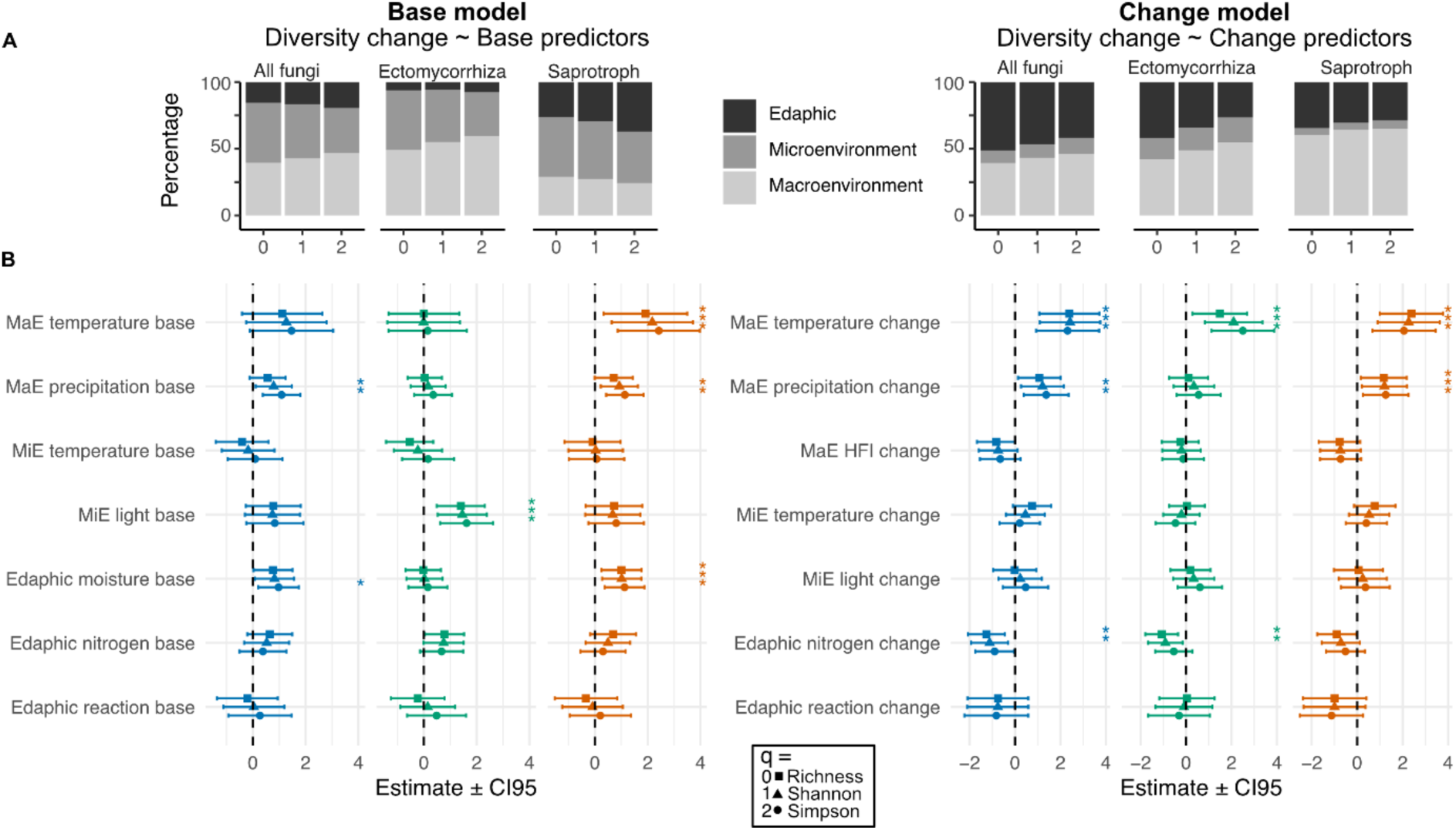
**Effects of base and change environmental variables on European fungal alpha diversity change**. Generalized additive models (GAM) with parametric predictors and a spatial smooth term. A) Hierarchical partitioning of explained variance by three predictor sets: Macroenvironmental (MaE), Microenvironmental (MiE) and edaphic. B) Multivariate GAM with detailed parametric (linear) predictor effects and spatial smooth term. For each variable Hill number = 0 is the top bar, and Hill number = 2 the bottom bar. Effects are shown as estimates with 95% confidence interval (CI). A star on the right next to the estimate indicates a significant effect (p<0.05). P values and thus significances were Bonferroni adjusted due to multiple testing by two models against the response variable. For the base and change models, the average deviance explained was 22.1% and 22.5%, respectively, with the inclusion of a spatial autocorrelation term.

We found a significantly positive relation of precipitation sum and moisture with diversity change for the Shannon (q=1) and Simpson (q=2) index for all fungi and saprotrophs. Considering the scatter plot and the partial regression line, the change was more pronounced at the dry end of the gradient for macroenvironmental precipitation and edaphic moisture, indicating that drought-prone areas experienced a loss of dominant species (Figs. 2, 3, S6). Ectomycorrhizal diversity change was significantly positively related to light (Figs. 2, 3). Considering the scatter plot and partial regression line (Fig. S6), the change was significant at both ends of the light gradient, indicating that grid cells characterized by shady conditions were associated with diversity loss and light conditions with diversity increase. We further found a significant positive relation between mean annual temperature and saprotrophic diversity change, indicating that colder climates experienced diversity loss while warmer areas experienced diversity gain (Fig. 2,3).

**Fig. 3.**
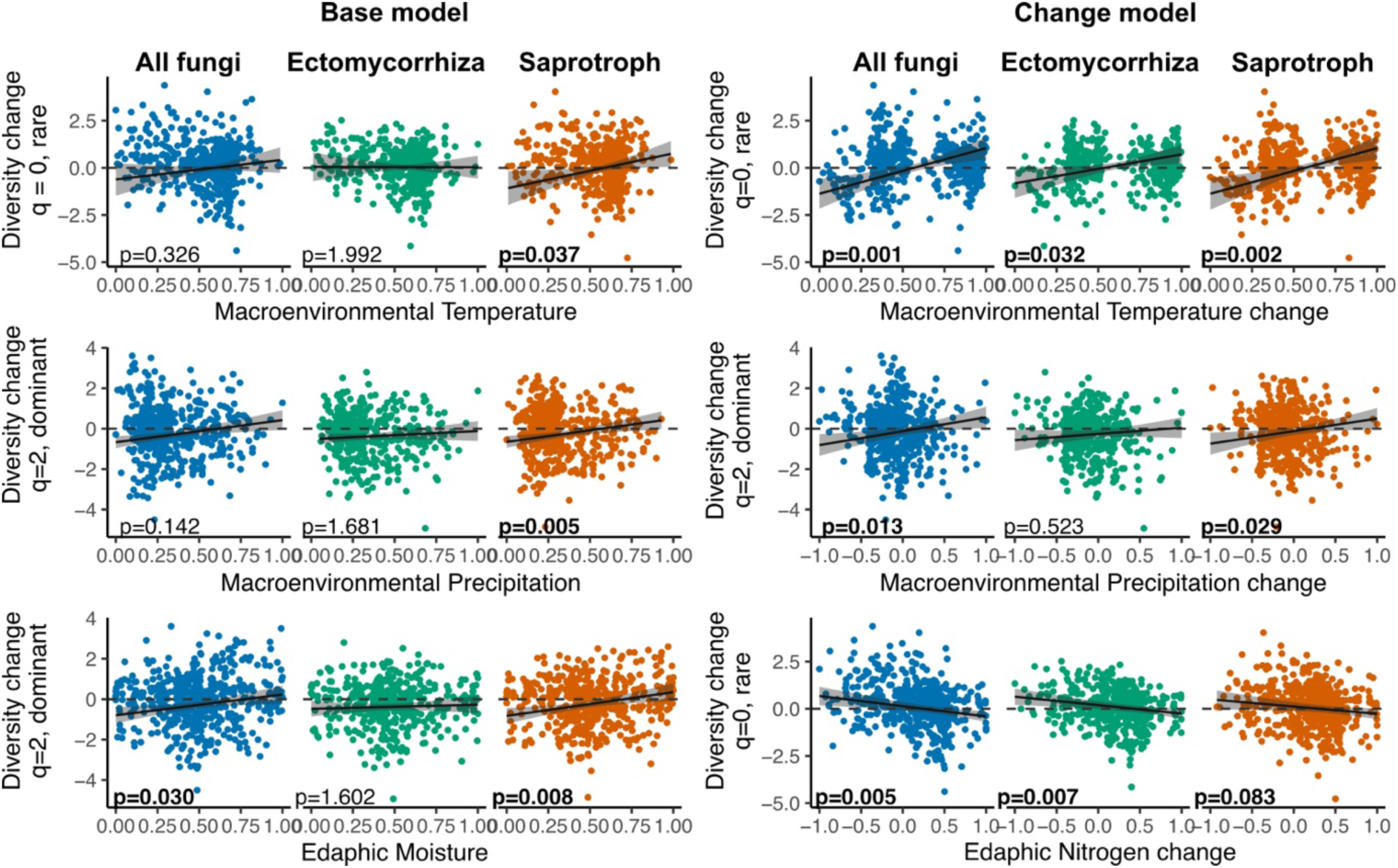
**The effects of selected base and change variables on changes in European fungal alpha diversity**. The lines represent the partial regression estimates from the multivariate linear model (Fig. 2, Tables S1,2). Moisture and nitrogen were estimated based on average plant Ellenberg indicator values. Nitrogen change is the residuals (linear model change relative to the base), where positive and negative values indicate an increase or decrease relative to the base. The x-axis was rescaled to range between 0 and 1 or-1 and 1, according to the original variable.

Within the change model, we found a significantly positive effect of mean annual temperature change on diversity change across all fungi and functional groups and the Hill series (Figs. 2,3, S7). Considering the scatter plot and partial regression line, the change was significant at both ends of the gradient, indicating an increase in diversity in areas that warmed strongly and a decrease in regions that warmed less strongly (Figs. 2 and 3). We did not add moisture as a predictor variable in the change model due to collinearity. We further found a significantly positive relationship between diversity change and precipitation change for all fungi and saprotrophs (Figs. 2, 3). Considering the scatter plot and partial regression line, the change was stronger at the lower end of the gradient, indicating a loss of diversity in areas that experienced decreased precipitation. We finally found a significant negative relationship between nitrogen change and diversity change (Figs. 2,3). This effect was significant for all fungi and ectomycorrhiza fungi. Based on the scatterplot and partial regression line, the change was more substantial at the negative end of the nitrogen gradient, indicating a gain in diversity where nitrogen (i.e., nitrogen input) has decreased (Figs. 2, 3). The trend was consistent for saprotrophs but not significant. For all scatter plots with univariate model statistics, see Figs. S6, 7.

## Discussion

Our analysis of a unique spatiotemporal dataset containing 5 million fungal records across Europe revealed only a marginal net change in alpha diversity over the last decades. This result aligns well with previous findings for other organism groups, considering the same biogeographical region classification ^3–5^, At the detailed level, grid cells associated with a decrease in alpha diversity were characterized by generally dry conditions (low precipitation and moisture) and increasing levels of nitrogen load. In contrast, increases in diversity were associated with stronger warming. Geographically, we found negative diversity changes in the Atlantic regions (Fig. S4). This negative diversity trend was stronger when common and dominant species were emphasized (Hill q>0). The Atlantic region is considered one of the most heavily modified in Europe, with the highest share of habitats assessed as in unfavourable or poor conservation status among EU biogeographical regions^4546474849^. In general, the neutral or even increasing effects across most parts of Europe might be due to different factors. Europe has undergone repeated waves of land-use intensification, deforestation, and habitat transformation long before the industrial era ^50,51^. Looking at the post-1970 period, our results suggest that fungal community-level diversity is, on average, relatively tolerant to ongoing global change drivers. Nevertheless, distinct regions show exceptionally strong positive or negative responses, and our dataset provides the resolution needed to identify and interpret these contrasting patterns.

Across functional groups, we observed an increase in diversity with macroclimatic warming, especially in warmer areas, suggesting that growth and fruiting conditions have become more favorable for many species. The result is in contrast to previous spatial analyses of fungal diversity, which have shown a hump-shaped effect of mean annual temperature on fungal alpha diversity both from metabarcoding and sporocarp data ^12,22^. This contrast highlights the importance of temporal data, as space-for-time analyses seem to overestimate the relevance and direction of temperature. The positive association between warming and increased alpha diversity could be explained by either increased diversity in line with the species-energy hypothesis ^52^ and metabolic theory ^53^ or by an intensification of cues that trigger fruiting ^54^.

Interestingly, we found consistently a decrease in diversity linked to low precipitation, low moisture, and decreased precipitation, suggesting negative effects of drying regions (Fig. 3). Several studies based on metabarcoding rather than sporocarp data, have suggested drought tolerance to be prevalent in fungi within dry areas ^55,56^, where species likely had enough evolutionary time to adapt. Across Europe, fungal diversity was shown to be mostly tolerant to drought manipulations ^23^. We found stronger effects of precipitation and moisture-based change on dominant than rare fungal species (Fig. 2), suggesting that drought is negatively affecting especially the dominant taxa, which might not be able to tolerate dry conditions. This pattern can be explained well by the dominance-tolerance trade-off ^57^, stating that species are either dominant (in terms of competition) or tolerant. Thus, dominant taxa might be intolerant to increasing droughts, while rare or specialized taxa more tolerant. Another explanation is Grime’s C–S axis ^58^, where dominant species resemble competitors that thrive under benign conditions but are drought-intolerant. In this concept, ruderals represent a strategy focused on rapid colonization after disturbance, which might belong to the rare or common species rather than the dominant.

Besides climate, all fungi and ectomycorrhizal fungi showed a decrease in diversity with increasing or stable nitrogen (Fig. 4, S6). A recent meta-analysis of soil fungal diversity found that nitrogen deposition was among the most important factors ^24^. Across Europe, high nitrogen deposition has repeatedly been linked particularly to declines in ectomycorrhizal fungi ^25,59,60^. However, atmospheric nitrogen inputs have declined since around 1990 in many parts of Central and Western Europe ^61–63^, but lasting effects may have been caused to fungal communities adapted to naturally nitrogen-poor conditions. Two studies have found that ectomycorrhizal communities may recover from previous nitrogen loads. However, recovery was partial and slow, but stronger in areas where total nitrogen loads were lower ^64,65^. Further, nitrogen-sensitive species were replaced by more nitrogen-tolerant species in the Netherlands ^65^, and thus, species composition has not recovered to the original conditions. In line with this, we found a decrease in ectomycorrhizal diversity linked to higher nitrogen increase. This pattern was significant for rare and common species but not when dominant species were emphasized (Fig. 2B). This suggests that dominant and likely nitrogen-tolerant taxa are increasing in areas where the vegetation data suggest lasting effects of eutrophication; while rare and common species are negatively affected, suggesting lasting negative effects of eutrophication on ectomycorrhizal diversity. Thus, our study suggests that the large-scale (nation-wide, e.g., EU Nitrates Directive (1991)) environmental measures to reduce nitrogen inputs were only partly successful, underscoring the need for further actions to mitigate pollution as a significant threat not only to fungal biodiversity, but also to ecosystem functioning.

In conclusion, we studied the change in fungal alpha diversity across large spatial scales using a unique spatiotemporal dataset and found no evidence of continental-scale vulnerability to climate warming. However, increasing drought and pollution (nitrogen) emerged as a greater threat than warming per se. In contrast to metabarcoding studies, assessments based on sporocarp data show that fungal biodiversity appears to be vulnerable to drought and nitrogen loads. Sporocarp records indicate the presence of species and their successful reproduction and thus our results provide a more sensitive indicator of species’ relative fitness and how it responds to environmental changes. Decline in fungal diversity appears to be marked in Atlantic areas, probably reflecting negative impacts of habitat loss and other anthropogenic factors in recent decades. Hence, fungal conservation should focus on improving conditions, especially in these regions. Further, our results suggest a need for increased efforts to buffer or mitigate climate change and chemical pollution. Despite the conservation-relevant loss of species, we found mostly stable communities, indicating stability of ecosystems in terms of fungi-driven nutrient and carbon cycling.

## Supporting information

Suppmat

## Acknoledgements

This research was funded by Biodiversa+, the European Biodiversity Partnership, in the context of the“FunDive: Monitoring and mapping fungal diversity for nature conservation” project under the 2022–2023 BioDivMon joint call. It was co-funded by the European Commission (grant agreement No. 2128-00020A - Biodiversa2022-640.

## Data availability

The full summary statistics to support the findings of this study, and data used to generate the main figures are included within the Supplementary Data file.

## Authors contributions

F-S.K. designed, conceptualized, analysed the study and drafted the manuscript.

M.Z. helped with data curation and formal analysis. M.Z., C.B. and J.H-C. provided intellectual input and contributed writing the final draft. All authors reviewed the manuscript and approved the submitted final manuscript.

