## Supplementary material for "Temporal changes in macrofungal alpha diversity over four decades in Europe": Suppmat

#### Supplementary Figures

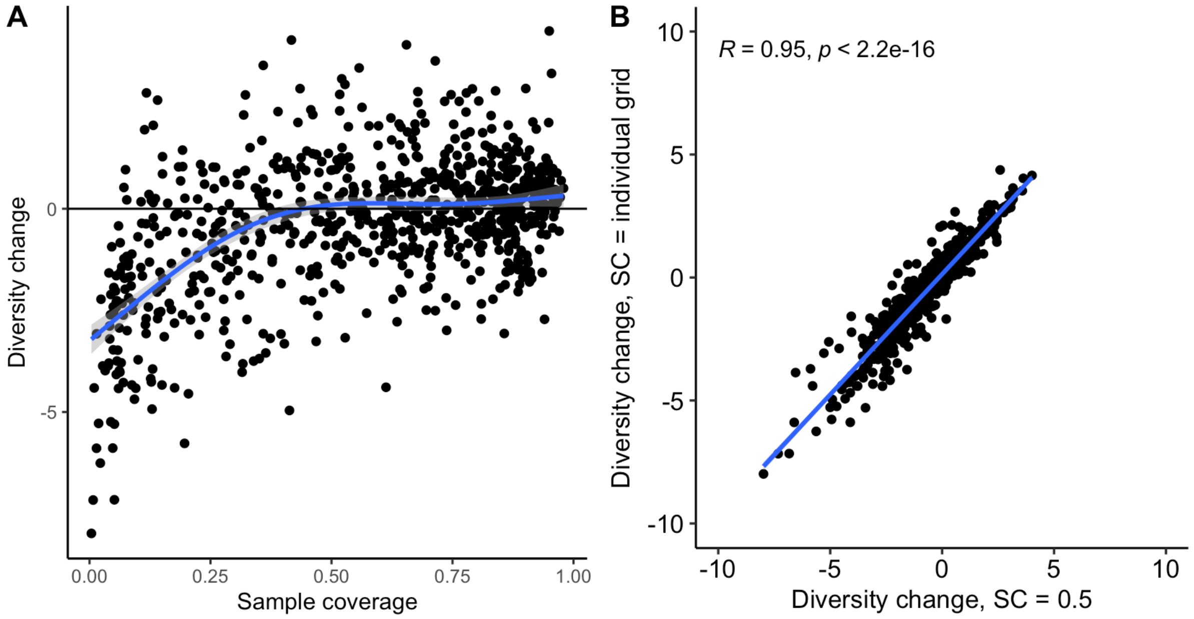


**Fig. S1 Diversity change in relation to the sample coverage and diversity change based on fixed sample coverage across grids**. A) The diversity change was biased towards negative values below a sample coverage of 0.5. We therefore used only grid cells with a minimum sample coverage of 0.5 for our analyses. B) We used for each grid separately the automatically chosen sample coverage for rarefaction and extrapolation. As we studied log ratios of diversity change, those are generally comparable between grid cells. However, differences in sample coverage might bias results. We therefore also used SC=0.5 as an absolute rarefaction and extrapolation level across grid cells in a second approach to calculate diversity change. We then correlated the diversity change of both approaches and found a strong and significant correlation between both variables. We thus continued with the diversity change values from the individual sample coverage approach as it maximises data use.

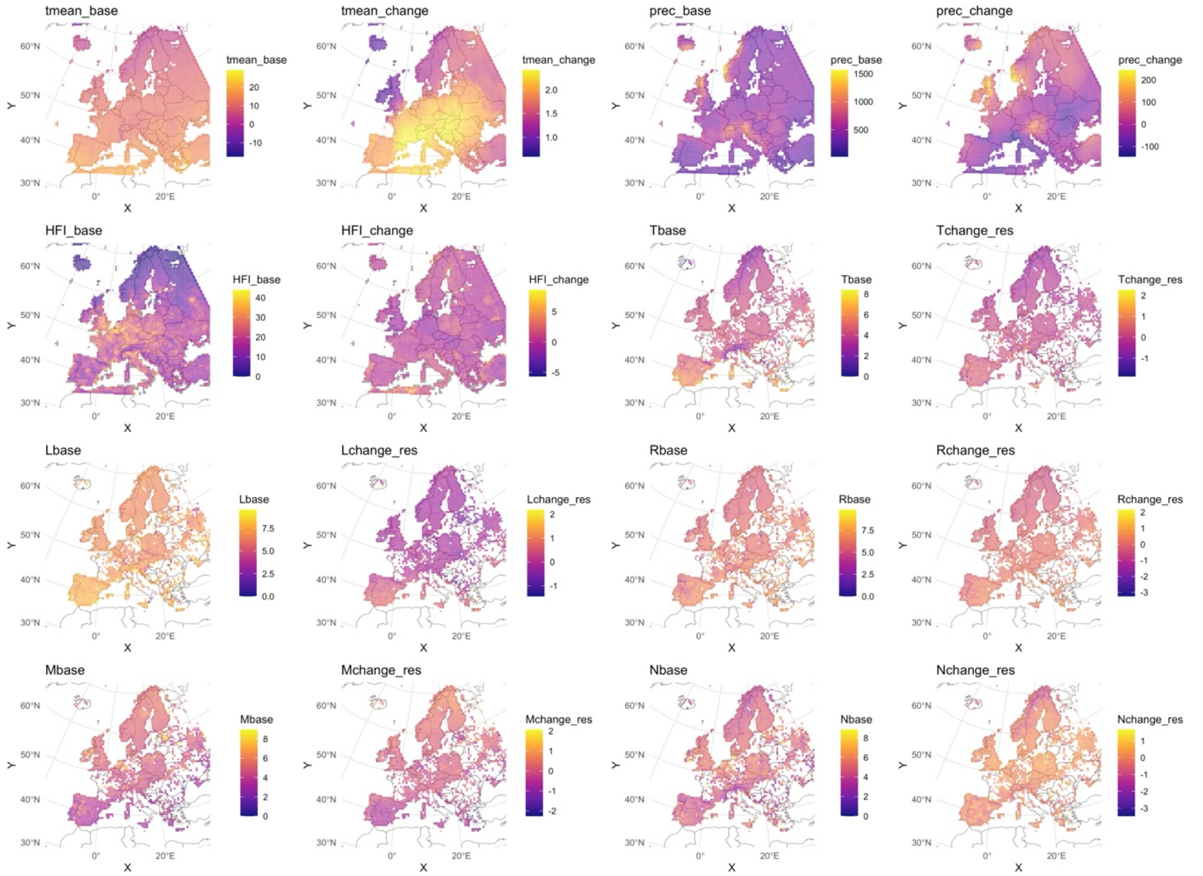


**Fig. S2 Maps of the predictor variables.** Macroenvironmental variables are tmean_base, tmean_change, prec_base, prec_change, HFI_base and HFI_change. Macroclimatic data have a base period of 1970-1990, and the change was calculated as the difference between 2010-2024 and the base. For the human footprint index (HFI), we used the first year (2000) as the base year, and the change was calculated as the difference between the last year (2018) and the base year. Microenvironmental and edaphic variables were based on the means of vegetation Ellenberg indicator values for each grid cell. Microenvironmental variables were Tbase, Tchange_res, Lbase, and Lchange_res. Edaphic variables were Mbase, Mchange_res, Rbase, R_change_res, Nbase and Nbase_res. The change values with "_res" are the residuals from linear models regressing "base" on the "change" variable to retrieve the change relative to the base values, as both were strongly correlated (Fig. S2).

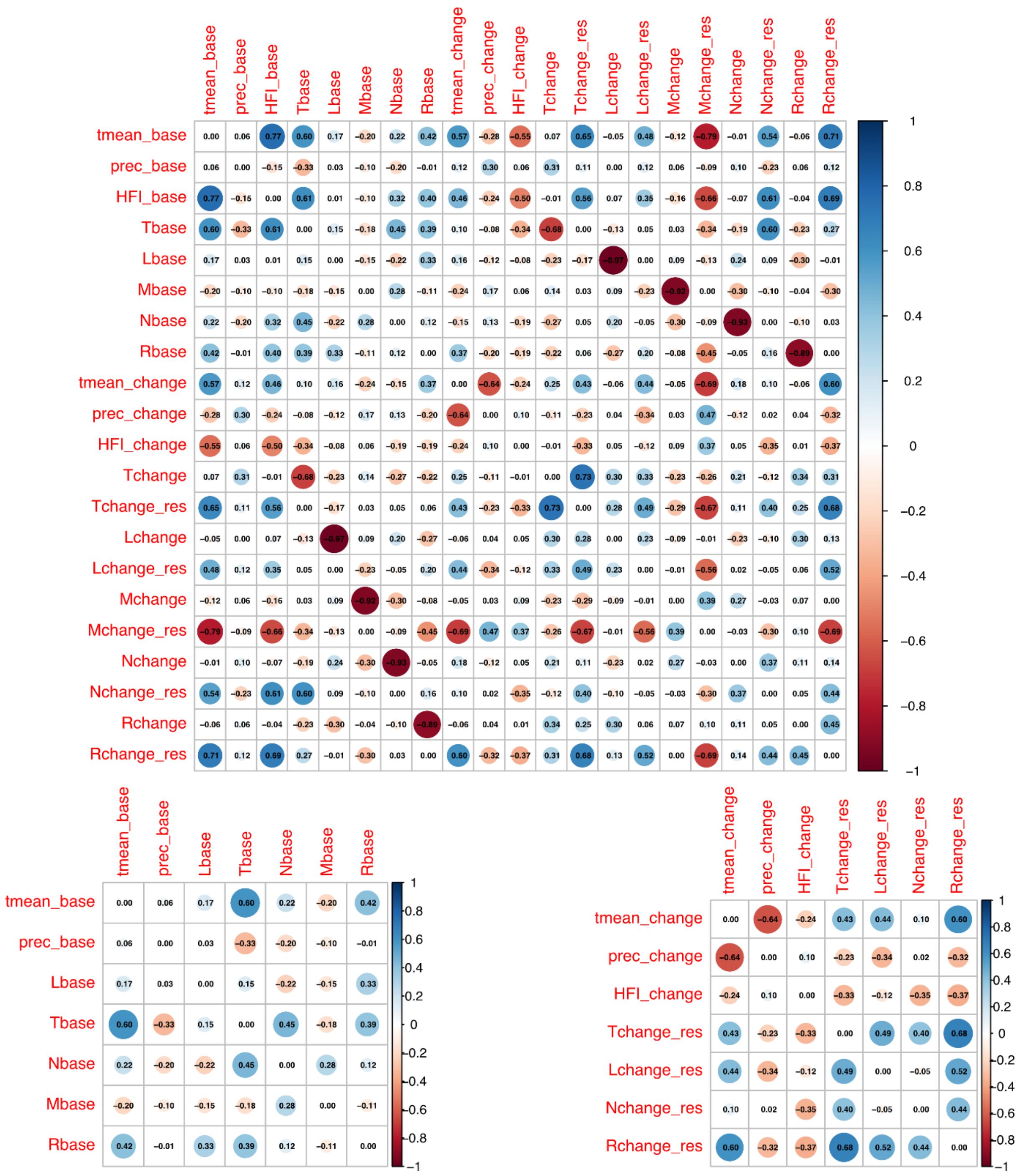


**Fig. S3 Correlation matrix of predictor values.** Top are all predictor values. Bottom are the selected base predictors (right) and change predictors (left).

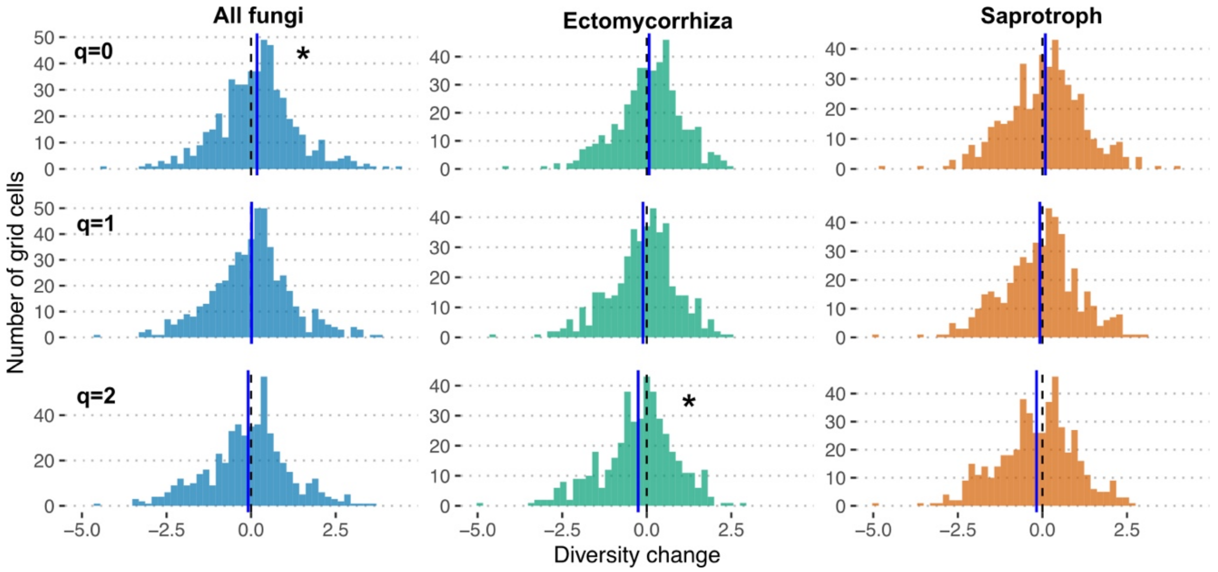


**Fig. S4** Alpha diversity is based on a coverage-based rarefaction and interpolation of incidence data per grid cell. The diversity change was calculated as log ratios between 1970-1990 and 2014-2024. Histograms of diversity change for alpha diversity. The zero point and change mean are drawn as dashed and solid lines, respectively. The mean was tested against zero using one parametric and two non-parametric tests, and we considered their significance (indicated by a star) only if all three tests showed a p-value of less than 0.05.

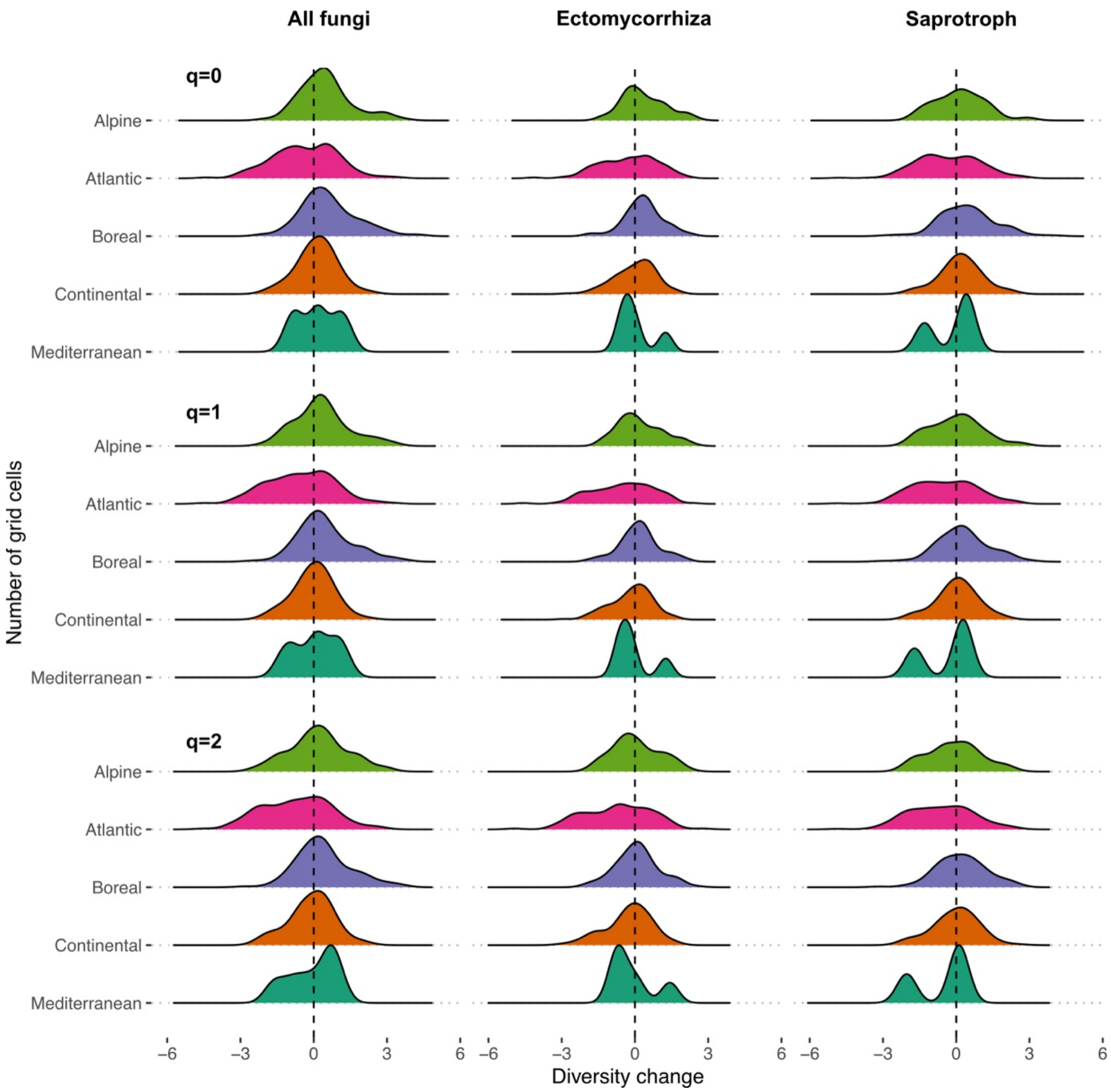


**Fig. S5 Exploration of alpha diversity change with European biogeographical regions**. Biogeographical regions were classified following [43]. The Arctic region was removed due to data insufficiency. The Mediterranean had also considerably less data than the other biogeographical regions. The Atlantic region showed more pronounced negative diversity changes, which is due to the UK showing more grid cells with negative changes (Fig. 1).

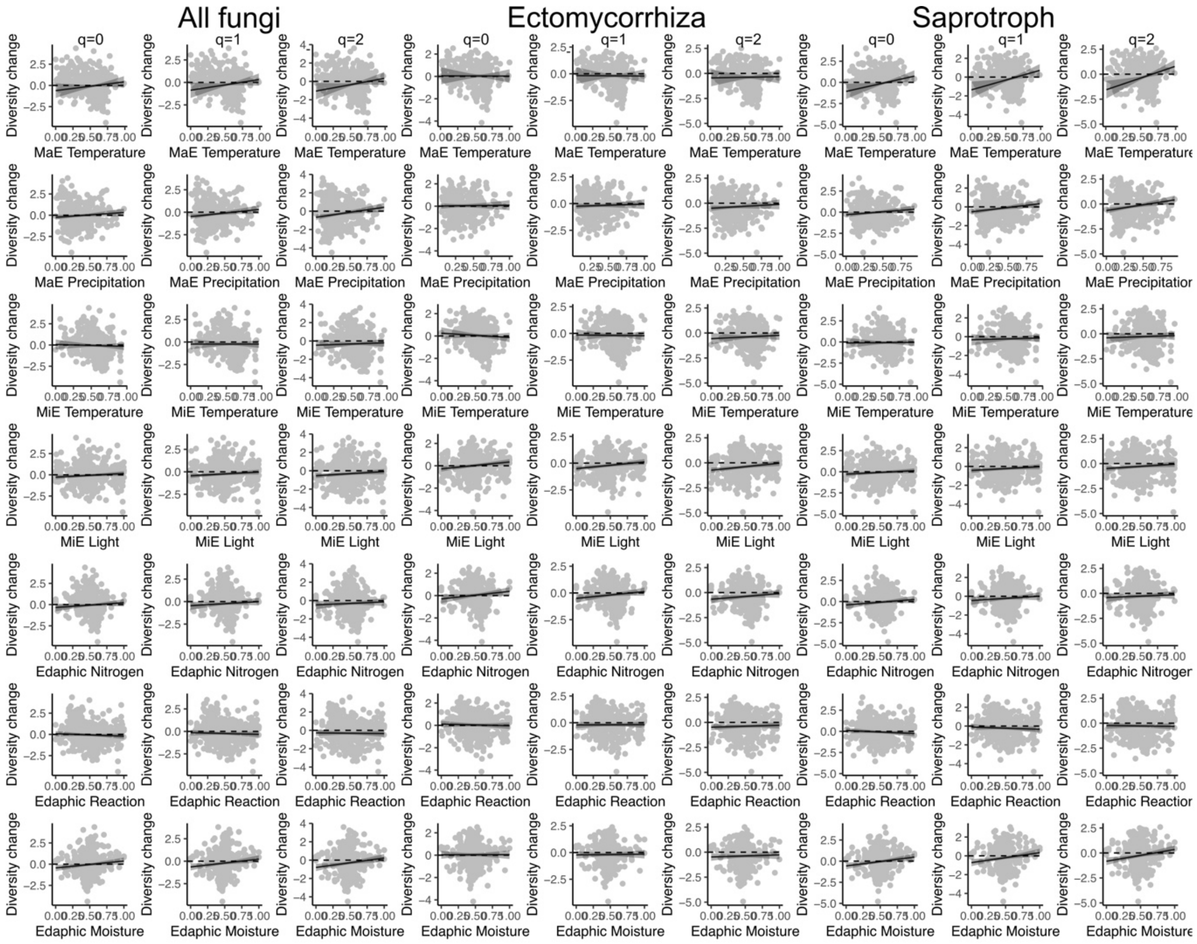


**Fig. S6 The effects of base variables on changes in European alpha diversity**. The lines show the partial estimates of the multivariate linear model predictions for all fungi, ectomycorrhizal and saprotrophic fungi (Fig. 2, Tables S1,2). The x-axis was rescaled between 0 and 1 or -1 and 1 accordingly.

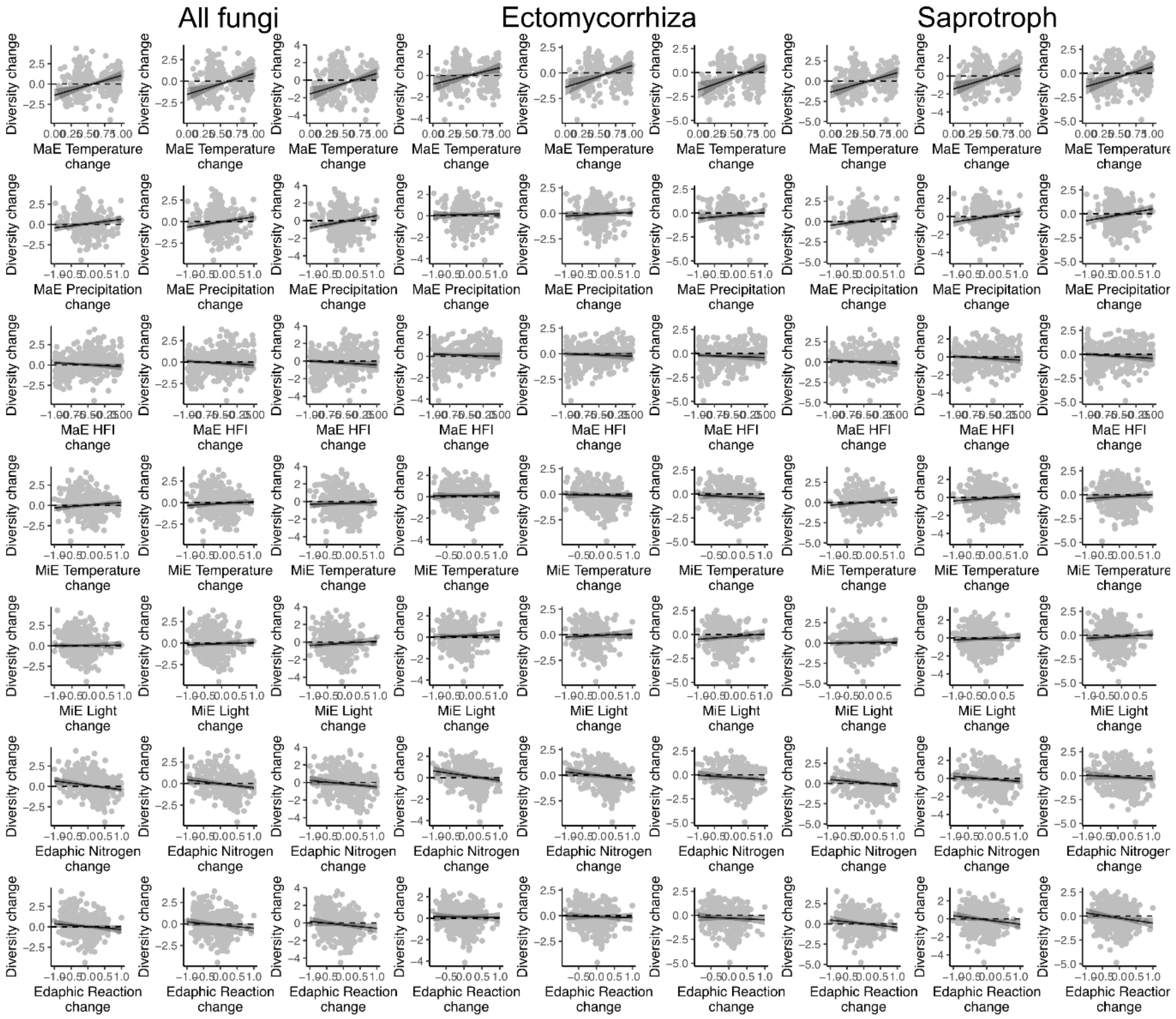


**Fig. S7 The effects of change variables on changes in European alpha diversity**. The lines show the partial estimates of the multivariate linear model predictions for all fungi, ectomycorrhizal and saprotrophic fungi (Fig. 2, Tables S1,2). The x-axis was rescaled between 0 and 1 or -1 and 1 accordingly.

### Supplementary Tables

**Table S1** Effects of the generalized additive model (GAM) with parametric (linear) environmental base predictors on alpha diversity change across Europe along the Hill series. The model contained a spatial correlation term. MaE = Macroenvironment; MiE = Microenvironment. Significant effects are highlighted in bold. t = t-value; p.adj = adjusted p value (Bonferroni); CI = 95% confidence interval.

| Group | Order.q | Predictor | Estimate | CI | StE | t | p adj. |
|---|---|---|---|---|---|---|---|
| All fungi | 0 | Intercept | -1,05 | -2.00--0.10 | 0,48 | **-2,17** | **0,061** |
| All fungi | 0 | MaE temperature base | 1,09 | -0.44- 2.61 | 0,78 | 1,40 | 0,326 |
| All fungi | 0 | MaE precipitation base | 0,55 | -0.13- 1.23 | 0,35 | 1,58 | 0,228 |
| All fungi | 0 | MiE temperature base | -0,32 | -1.30- 0.65 | 0,50 | -0,65 | 1,032 |
| All fungi | 0 | MiE light base | 0,42 | -0.01- 0.86 | 0,22 | 1,90 | 0,115 |
| All fungi | 0 | Edaphic moisture base | 0,46 | 0.02- 0.90 | 0,23 | 2,03 | 0,086 |
| All fungi | 0 | Edaphic nitrogen base | 0,68 | -0.13- 1.48 | 0,41 | 1,65 | 0,199 |
| All fungi | 0 | Edaphic reaction base | -0,28 | -0.84- 0.29 | 0,29 | -0,97 | 0,668 |
| Ectomycorrhiza | 0 | Intercept | -0,28 | -1.15-0.59 | 0,44 | -0,63 | 1,060 |
| Ectomycorrhiza | 0 | MaE temperature base | 0,00 | -1.34-1.35 | 0,68 | 0,01 | 1,992 |
| Ectomycorrhiza | 0 | MaE precipitation base | 0,06 | -0.59-0.71 | 0,33 | 0,18 | 1,708 |
| Ectomycorrhiza | 0 | MiE temperature base | -0,42 | -1.30-0.46 | 0,45 | -0,94 | 0,695 |
| Ectomycorrhiza | 0 | MiE light base | 0,61 | 0.22-1.00 | 0,20 | **3,03** | **0,005** |
| Ectomycorrhiza | 0 | Edaphic moisture base | -0,06 | -0.47-0.35 | 0,21 | -0,29 | 1,539 |
| Ectomycorrhiza | 0 | Edaphic nitrogen base | 0,70 | 0.00-1.40 | 0,36 | 1,97 | 0,100 |
| Ectomycorrhiza | 0 | Edaphic reaction base | -0,14 | -0.65-0.37 | 0,26 | -0,53 | 1,188 |
| Saprotroph | 0 | Intercept | -1,90 | -2.91--0.90 | 0,51 | **-3,71** | **0,000** |
| Saprotroph | 0 | MaE temperature base | 1,91 | 0.33- 3.50 | 0,81 | **2,36** | **0,037** |
| Saprotroph | 0 | MaE precipitation base | 0,70 | -0.02- 1.42 | 0,37 | 1,92 | 0,112 |
| Saprotroph | 0 | MiE temperature base | -0,03 | -1.07- 1.01 | 0,53 | -0,06 | 1,905 |
| Saprotroph | 0 | MiE light base | 0,41 | -0.05- 0.86 | 0,23 | 1,75 | 0,161 |
| Saprotroph | 0 | Edaphic moisture base | 0,60 | 0.15- 1.06 | 0,23 | **2,59** | **0,020** |
| Saprotroph | 0 | Edaphic nitrogen base | 0,79 | -0.03- 1.61 | 0,42 | 1,89 | 0,118 |
| Saprotroph | 0 | Edaphic reaction base | -0,34 | -0.93- 0.25 | 0,30 | -1,14 | 0,512 |
| All fungi | 1 | Intercept | -1,51 | -2.46--0.56 | 0,49 | **-3,12** | **0,004** |
| All fungi | 1 | MaE temperature base | 1,25 | -0.27- 2.78 | 0,78 | 1,61 | 0,217 |
| All fungi | 1 | MaE precipitation base | 0,79 | 0.10- 1.47 | 0,35 | **2,26** | **0,049** |
| All fungi | 1 | MiE temperature base | -0,10 | -1.08- 0.88 | 0,50 | -0,20 | 1,676 |
| All fungi | 1 | MiE light base | 0,42 | -0.02- 0.86 | 0,22 | 1,88 | 0,122 |
| All fungi | 1 | Edaphic moisture base | 0,49 | 0.05- 0.94 | 0,23 | 2,17 | 0,060 |
| All fungi | 1 | Edaphic nitrogen base | 0,58 | -0.23- 1.38 | 0,41 | 1,41 | 0,319 |
| All fungi | 1 | Edaphic reaction base | -0,17 | -0.74- 0.39 | 0,29 | -0,59 | 1,107 |
| Ectomycorrhiza | 1 | Intercept | -0,77 | -1.66-0.12 | 0,46 | **-1,69** | **0,185** |
| Ectomycorrhiza | 1 | MaE temperature base | -0,01 | -1.39-1.37 | 0,70 | -0,02 | 1,974 |
| Ectomycorrhiza | 1 | MaE precipitation base | 0,19 | -0.47-0.86 | 0,34 | 0,57 | 1,133 |
| Ectomycorrhiza | 1 | MiE temperature base | -0,11 | -1.01-0.79 | 0,46 | -0,24 | 1,622 |
| Ectomycorrhiza | 1 | MiE light base | 0,63 | 0.23-1.04 | 0,21 | **3,09** | **0,004** |
| Ectomycorrhiza | 1 | Edaphic moisture base | -0,02 | -0.44-0.39 | 0,21 | -0,11 | 1,820 |

| | | | | | | | |
|---|---|---|---|---|---|---|---|
| Ectomycorrhiza | 1 | Edaphic nitrogen base | 0,67 | -0.04-1.39 | 0,36 | 1,85 | 0,131 |
| Ectomycorrhiza | 1 | Edaphic reaction base | 0,03 | -0.49-0.55 | 0,27 | 0,12 | 1,801 |
| Saprotroph | 1 | Intercept | -2,30 | -3.28--1.32 | 0,50 | **-4,60** | **0,000** |
| Saprotroph | 1 | MaE temperature base | 2,18 | 0.63- 3.72 | 0,79 | **2,76** | **0,012** |
| Saprotroph | 1 | MaE precipitation base | 0,92 | 0.22- 1.62 | 0,36 | **2,58** | **0,020** |
| Saprotroph | 1 | MiE temperature base | 0,08 | -0.94- 1.09 | 0,52 | 0,15 | 1,769 |
| Saprotroph | 1 | MiE light base | 0,39 | -0.05- 0.84 | 0,23 | 1,75 | 0,161 |
| Saprotroph | 1 | Edaphic moisture base | 0,60 | 0.16- 1.05 | 0,23 | **2,66** | **0,016** |
| Saprotroph | 1 | Edaphic nitrogen base | 0,61 | -0.19- 1.41 | 0,41 | 1,50 | 0,268 |
| Saprotroph | 1 | Edaphic reaction base | -0,22 | -0.79- 0.35 | 0,29 | -0,76 | 0,895 |
| All fungi | 2 | Intercept | -2,02 | -3.01--1.03 | 0,50 | **-4,01** | **0,000** |
| All fungi | 2 | MaE temperature base | 1,46 | -0.12- 3.04 | 0,81 | 1,81 | 0,143 |
| All fungi | 2 | MaE precipitation base | 1,09 | 0.38- 1.80 | 0,36 | **3,01** | **0,006** |
| All fungi | 2 | MiE temperature base | 0,14 | -0.87- 1.16 | 0,52 | 0,28 | 1,560 |
| All fungi | 2 | MiE light base | 0,45 | 0.00- 0.91 | 0,23 | 1,95 | 0,103 |
| All fungi | 2 | Edaphic moisture base | 0,57 | 0.11- 1.03 | 0,24 | **2,44** | **0,030** |
| All fungi | 2 | Edaphic nitrogen base | 0,47 | -0.36- 1.31 | 0,43 | 1,11 | 0,535 |
| All fungi | 2 | Edaphic reaction base | -0,06 | -0.65- 0.53 | 0,30 | -0,20 | 1,681 |
| Ectomycorrhiza | 2 | Intercept | -1,40 | -2.37--0.44 | 0,49 | **-2,84** | **0,009** |
| Ectomycorrhiza | 2 | MaE temperature base | 0,15 | -1.34- 1.64 | 0,76 | 0,20 | 1,681 |
| Ectomycorrhiza | 2 | MaE precipitation base | 0,39 | -0.32- 1.11 | 0,36 | 1,08 | 0,562 |
| Ectomycorrhiza | 2 | MiE temperature base | 0,27 | -0.70- 1.24 | 0,49 | 0,55 | 1,163 |
| Ectomycorrhiza | 2 | MiE light base | 0,70 | 0.26- 1.13 | 0,22 | **3,15** | **0,003** |
| Ectomycorrhiza | 2 | Edaphic moisture base | 0,06 | -0.39- 0.51 | 0,23 | 0,25 | 1,602 |
| Ectomycorrhiza | 2 | Edaphic nitrogen base | 0,62 | -0.15- 1.39 | 0,39 | 1,57 | 0,235 |
| Ectomycorrhiza | 2 | Edaphic reaction base | 0,20 | -0.36- 0.76 | 0,29 | 0,69 | 0,986 |
| Saprotroph | 2 | Intercept | -2,66 | -3.65--1.67 | 0,51 | **-5,27** | **0,000** |
| Saprotroph | 2 | MaE temperature base | 2,44 | 0.88- 4.00 | 0,80 | **3,07** | **0,005** |
| Saprotroph | 2 | MaE precipitation base | 1,14 | 0.43- 1.84 | 0,36 | **3,16** | **0,003** |
| Saprotroph | 2 | MiE temperature base | 0,09 | -0.93- 1.12 | 0,52 | 0,18 | 1,714 |
| Saprotroph | 2 | MiE light base | 0,43 | -0.01- 0.88 | 0,23 | 1,90 | 0,117 |
| Saprotroph | 2 | Edaphic moisture base | 0,66 | 0.21- 1.11 | 0,23 | **2,88** | **0,008** |
| Saprotroph | 2 | Edaphic nitrogen base | 0,44 | -0.36- 1.25 | 0,41 | 1,07 | 0,568 |
| Saprotroph | 2 | Edaphic reaction base | -0,07 | -0.65- 0.51 | 0,29 | -0,24 | 1,628 |

**Table S2** Effects of the generalized additive model (GAM) with parametric (linear) environmental change predictors on alpha diversity change across Europe along the Hill series. The model contained a spatial correlation term. MaE = Macroenvironment; MiE = Microenvironment. Significant effects are highlighted in bold. t = t-value; p.adj = adjusted p value (Bonferroni); CI = 95% confidence interval.

| Group | Order.q | Predictor | Estimate | CI | StE | t | p |
|---|---|---|---|---|---|---|---|
| All fungi | 0 | Intercept | -1,42 | -2.25--0.58 | 0,43 | **-3,33** | **0,002** |
| All fungi | 0 | MaE temperature change | 2,42 | 1.10- 3.74 | 0,67 | **3,58** | **0,001** |
| All fungi | 0 | MaE precipitation change | 0,52 | 0.06- 0.99 | 0,24 | 2,21 | 0,055 |
| All fungi | 0 | MaE HFI change | -0,49 | -0.95--0.03 | 0,24 | -2,08 | 0,076 |
| All fungi | 0 | MiE temperature change | 0,33 | -0.04- 0.70 | 0,19 | 1,76 | 0,159 |
| All fungi | 0 | MiE light change | 0,04 | -0.37- 0.46 | 0,21 | 0,20 | 1,679 |
| All fungi | 0 | Edaphic nitrogen change | -0,54 | -0.88--0.21 | 0,17 | **-3,16** | **0,003** |
| All fungi | 0 | Edaphic reaction change | -0,35 | -0.84- 0.14 | 0,25 | -1,38 | 0,335 |
| Ectomycorrhiza | 0 | Intercept | -0,89 | -1.68--0.10 | 0,40 | -2,21 | 0,055 |
| Ectomycorrhiza | 0 | MaE temperature change | 1,54 | 0.33- 2.75 | 0,62 | **2,49** | **0,026** |
| Ectomycorrhiza | 0 | MaE precipitation change | 0,07 | -0.36- 0.49 | 0,22 | 0,31 | 1,510 |
| Ectomycorrhiza | 0 | MaE HFI change | -0,21 | -0.65- 0.22 | 0,22 | -0,96 | 0,678 |
| Ectomycorrhiza | 0 | MiE temperature change | 0,03 | -0.31- 0.37 | 0,17 | 0,17 | 1,735 |
| Ectomycorrhiza | 0 | MiE light change | 0,10 | -0.29- 0.49 | 0,20 | 0,51 | 1,215 |
| Ectomycorrhiza | 0 | Edaphic nitrogen change | -0,45 | -0.75--0.15 | 0,15 | **-2,94** | **0,007** |
| Ectomycorrhiza | 0 | Edaphic reaction change | -0,04 | -0.48- 0.41 | 0,23 | -0,16 | 1,739 |
| Saprotroph | 0 | Intercept | -1,49 | -2.39--0.60 | 0,46 | **-3,28** | **0,002** |
| Saprotroph | 0 | MaE temperature change | 2,41 | 1.01- 3.81 | 0,71 | **3,38** | **0,002** |
| Saprotroph | 0 | MaE precipitation change | 0,57 | 0.07- 1.07 | 0,26 | **2,23** | **0,053** |
| Saprotroph | 0 | MaE HFI change | -0,44 | -0.93- 0.05 | 0,25 | -1,76 | 0,160 |
| Saprotroph | 0 | MiE temperature change | 0,36 | -0.04- 0.76 | 0,20 | 1,78 | 0,152 |
| Saprotroph | 0 | MiE light change | 0,07 | -0.40- 0.53 | 0,24 | 0,28 | 1,558 |
| Saprotroph | 0 | Edaphic nitrogen change | -0,38 | -0.74--0.03 | 0,18 | -2,12 | 0,070 |
| Saprotroph | 0 | Edaphic reaction change | -0,45 | -0.96- 0.06 | 0,26 | -1,72 | 0,172 |
| All fungi | 1 | Intercept | -1,55 | -2.39--0.71 | 0,43 | **-3,62** | **0,001** |
| All fungi | 1 | MaE temperature change | 2,44 | 1.11- 3.77 | 0,68 | **3,61** | **0,001** |
| All fungi | 1 | MaE precipitation change | 0,60 | 0.13- 1.06 | 0,24 | **2,50** | **0,025** |
| All fungi | 1 | MaE HFI change | -0,45 | -0.92- 0.01 | 0,24 | -1,91 | 0,113 |
| All fungi | 1 | MiE temperature change | 0,21 | -0.16- 0.59 | 0,19 | 1,12 | 0,528 |
| All fungi | 1 | MiE light change | 0,15 | -0.26- 0.57 | 0,21 | 0,72 | 0,943 |
| All fungi | 1 | Edaphic nitrogen change | -0,47 | -0.81--0.13 | 0,17 | **-2,72** | **0,013** |
| All fungi | 1 | Edaphic reaction change | -0,37 | -0.86- 0.13 | 0,25 | -1,46 | 0,289 |
| Ectomycorrhiza | 1 | Intercept | -1,42 | -2.25--0.59 | 0,42 | **-3,36** | **0,002** |
| Ectomycorrhiza | 1 | MaE temperature change | 2,13 | 0.86- 3.40 | 0,65 | **3,29** | **0,002** |
| Ectomycorrhiza | 1 | MaE precipitation change | 0,19 | -0.25- 0.63 | 0,23 | 0,85 | 0,797 |
| Ectomycorrhiza | 1 | MaE HFI change | -0,22 | -0.68- 0.23 | 0,23 | -0,96 | 0,676 |
| Ectomycorrhiza | 1 | MiE temperature change | -0,07 | -0.43- 0.28 | 0,18 | -0,42 | 1,351 |

| | | | | | | | |
|---|---|---|---|---|---|---|---|
| Ectomycorrhiza | 1 | MiE light change | 0,16 | -0.24- 0.56 | 0,20 | 0,79 | 0,863 |
| Ectomycorrhiza | 1 | Edaphic nitrogen change | -0,37 | -0.68--0.06 | 0,16 | **-2,32** | **0,042** |
| Ectomycorrhiza | 1 | Edaphic reaction change | -0,08 | -0.54- 0.37 | 0,23 | -0,36 | 1,440 |
| Saprotroph | 1 | Intercept | -1,56 | -2.43--0.68 | 0,44 | **-3,50** | **0,001** |
| Saprotroph | 1 | MaE temperature change | 2,28 | 0.92- 3.65 | 0,70 | **3,27** | **0,002** |
| Saprotroph | 1 | MaE precipitation change | 0,59 | 0.10- 1.08 | 0,25 | **2,34** | **0,039** |
| Saprotroph | 1 | MaE HFI change | -0,42 | -0.90- 0.06 | 0,24 | -1,72 | 0,174 |
| Saprotroph | 1 | MiE temperature change | 0,26 | -0.13- 0.65 | 0,20 | 1,31 | 0,379 |
| Saprotroph | 1 | MiE light change | 0,15 | -0.30- 0.61 | 0,23 | 0,66 | 1,016 |
| Saprotroph | 1 | Edaphic nitrogen change | -0,30 | -0.64- 0.05 | 0,18 | -1,68 | 0,188 |
| Saprotroph | 1 | Edaphic reaction change | -0,47 | -0.97- 0.03 | 0,26 | -1,85 | 0,129 |
| All fungi | 2 | Intercept | -1,56 | -2.44--0.68 | 0,45 | **-3,48** | **0,001** |
| All fungi | 2 | MaE temperature change | 2,34 | 0.95- 3.73 | 0,71 | **3,29** | **0,002** |
| All fungi | 2 | MaE precipitation change | 0,69 | 0.20- 1.18 | 0,25 | **2,74** | **0,013** |
| All fungi | 2 | MaE HFI change | -0,43 | -0.91- 0.06 | 0,25 | -1,73 | 0,168 |
| All fungi | 2 | MiE temperature change | 0,11 | -0.28- 0.50 | 0,20 | 0,56 | 1,153 |
| All fungi | 2 | MiE light change | 0,26 | -0.18- 0.69 | 0,22 | 1,16 | 0,496 |
| All fungi | 2 | Edaphic nitrogen change | -0,36 | -0.72--0.01 | 0,18 | -2,01 | 0,089 |
| All fungi | 2 | Edaphic reaction change | -0,40 | -0.92- 0.11 | 0,26 | -1,54 | 0,251 |
| Ectomycorrhiza | 2 | Intercept | -1,80 | -2.70--0.90 | 0,46 | **-3,92** | **0,000** |
| Ectomycorrhiza | 2 | MaE temperature change | 2,54 | 1.16- 3.92 | 0,70 | **3,61** | **0,001** |
| Ectomycorrhiza | 2 | MaE precipitation change | 0,31 | -0.17- 0.79 | 0,24 | 1,25 | 0,424 |
| Ectomycorrhiza | 2 | MaE HFI change | -0,22 | -0.72- 0.27 | 0,25 | -0,88 | 0,756 |
| Ectomycorrhiza | 2 | MiE temperature change | -0,19 | -0.57- 0.19 | 0,19 | -0,96 | 0,679 |
| Ectomycorrhiza | 2 | MiE light change | 0,28 | -0.15- 0.71 | 0,22 | 1,26 | 0,417 |
| Ectomycorrhiza | 2 | Edaphic nitrogen change | -0,21 | -0.55- 0.13 | 0,17 | -1,22 | 0,450 |
| Ectomycorrhiza | 2 | Edaphic reaction change | -0,16 | -0.65- 0.33 | 0,25 | -0,63 | 1,058 |
| Saprotroph | 2 | Intercept | -1,54 | -2.43--0.65 | 0,45 | **-3,40** | **0,001** |
| Saprotroph | 2 | MaE temperature change | 2,08 | 0.69- 3.47 | 0,71 | **2,93** | **0,007** |
| Saprotroph | 2 | MaE precipitation change | 0,62 | 0.12- 1.12 | 0,25 | **2,43** | **0,031** |
| Saprotroph | 2 | MaE HFI change | -0,43 | -0.92- 0.06 | 0,25 | -1,73 | 0,170 |
| Saprotroph | 2 | MiE temperature change | 0,21 | -0.18- 0.60 | 0,20 | 1,04 | 0,599 |
| Saprotroph | 2 | MiE light change | 0,21 | -0.26- 0.67 | 0,24 | 0,88 | 0,757 |
| Saprotroph | 2 | Edaphic nitrogen change | -0,20 | -0.56- 0.15 | 0,18 | -1,14 | 0,511 |
| Saprotroph | 2 | Edaphic reaction change | -0,53 | -1.04--0.02 | 0,26 | -2,05 | 0,082 |
